# A minimal synaptic model for direction selective neurons in *Drosophila*

**DOI:** 10.1101/833970

**Authors:** Jacob A. Zavatone-Veth, Bara A. Badwan, Damon A. Clark

**Affiliations:** Department of Physics, Yale University, New Haven, CT 06511, USA; Department of Molecular, Cellular, and Developmental Biology, Yale University, New Haven, CT 06511, USA; School of Engineering and Applied Science, Yale University, New Haven, CT 06511, USA; Interdepartmental Neuroscience Program, Yale University, New Haven, CT 06511, USA; Department of Neuroscience, Yale University, New Haven, CT 06511, USA

## Abstract

Visual motion estimation is a canonical neural computation. In *Drosophila*, recent advances have identified anatomical and functional circuitry underlying direction-selective computations. Models with varying levels of abstraction have been proposed to explain specific experimental results, but have rarely been compared across experiments. Here we construct a minimal, biophysically inspired synaptic model for *Drosophila*’s first-order direction-selective T4 cells using the wealth of available anatomical and physiological data. We show how this model relates mathematically to classical models of motion detection, including the Hassenstein-Reichardt correlator model. We used numerical simulation to test how well this synaptic model could reproduce measurements of T4 cells across many datasets and stimulus modalities. These comparisons include responses to sinusoid gratings, to apparent motion stimuli, to stochastic stimuli, and to natural scenes. Without fine-tuning this model, it sufficed to reproduce many, but not all, response properties of T4 cells. Since this model is flexible and based on straightforward biophysical properties, it provides an extensible framework for developing a mechanistic understanding of T4 neural response properties. Moreover, it can be used to assess the sufficiency of simple biophysical mechanisms to describe features of the direction-selective computation and identify where our understanding must be improved.

## Introduction

Motion estimation is a canonical visual computation that requires integrating information nonlinearly over both time and space. Direction-selective signals are tuned to motion in a preferred-direction (PD), which elicits the strongest responses, while motion in the opposite, null-direction (ND), elicits a weaker response. This directional computation has been described by a wide variety of computational models. Classical models, such as the Hassenstein-Reichardt correlator (HRC) (Hassenstein and Reichardt, 1956) and motion energy model (Adelson and Bergen, 1985), rely on sensing correlations between pairs points separated in time and space. These phenomenological models have provided striking insights into neural and behavioral responses in a variety of species, including in flies (Yang and Clandinin, 2018).

In the last decade, advances in defining the anatomical and functional connectivity of *Drosophila*’s visual circuits suggest that we should move towards more mechanistic, biophysical descriptions of this computation. Here, we follow previous work (Gruntman et al., 2018; Torre and Poggio, 1978) to propose a simple, biophysically-plausible synaptic model for direction-selectivity in Drosophila’s ON-edge sensitive motion pathway. We compare its predictions to measurements made by several research groups in response to many stimuli, giving us a tool for understanding which features are sufficient to describe different response properties.

The inputs to direction-selective cells have been identified by electron microscopy and through genetic silencing experiments. The most peripheral direction-selective neurons in the *Drosophila* optic lobe are the T4 and T5 cells, which are sensitive to moving ON-edges (consisting of contrast increments) and OFF-edges (consisting of contrast decrements), respectively (Clark et al., 2011; Joesch et al., 2010; Maisak et al., 2013). Electron microscopy and genetic silencing have identified primary inputs to T4 and T5 cells (Serbe et al., 2016; Shinomiya et al., 2019; Strother et al., 2017; Takemura et al., 2017). These studies suggest that T4 cells receive input from three distinct colinear spatial locations, with the neurons Mi1 and Tm3 both relaying information about the central point, and the neurons Mi9 and Mi4 acting as relays for the two flanking points (Takemura et al., 2017) (**Fig. 1A**). The neuron T5 appears to have a similar spatial structure, with different input neurons (Shinomiya et al., 2019). Both cell types also receive spatially-localized inputs from other neurons, whose functions remain less well understood (Shinomiya et al., 2019; Takemura et al., 2017).

**Figure 1:**
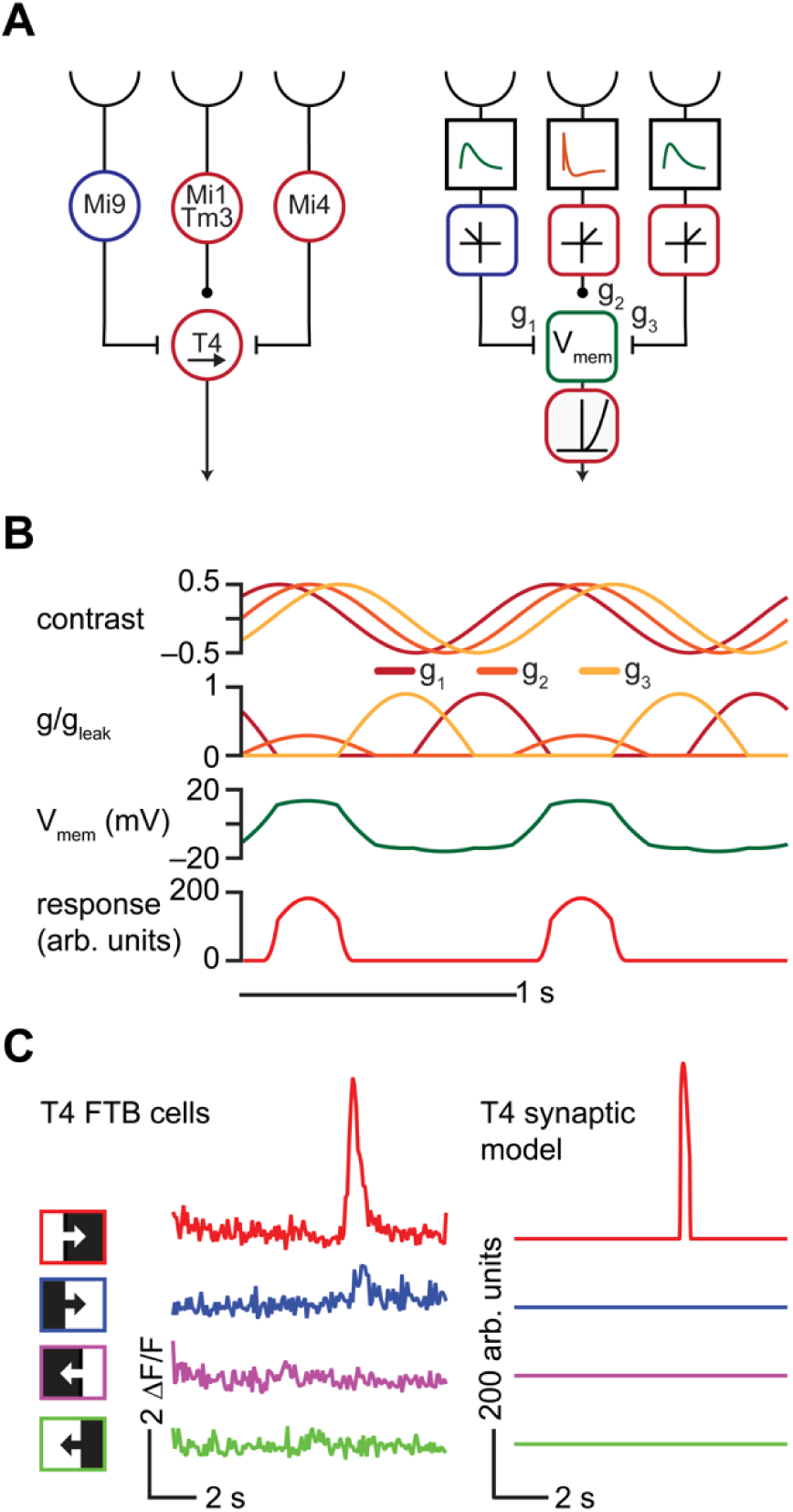
An anatomically constrained synaptic model for T4 cells. A. *Left:* Diagram of proposed inputs to *Drosophila* T4 first-order direction-selective cells based on anatomical and physiological measurements. Mi1 and Tm3 cells provide ON excitatory input at the center of the receptive field of each T4 cell, while Mi9 provides delayed OFF inhibitory input offset in the null direction, and Mi4 provides delayed ON inhibitory input offset in the preferred direction. *Right:* Synaptic model based on the anatomical structure shown at left. B. Responses of each component of the synaptic model to a 1 Hz, 45° sinusoidal grating drifting in the preferred (rightward) direction. *Top:* Input contrasts to each of the three presynaptic units of the model. *Upper middle:* Conductances of excitatory and inhibitory currents corresponding to each input in response to the sinusoidal stimulus. *Lower middle:* Membrane voltage. *Bottom:* Calcium signal. C. *Left:* Responses of T4 cells sensitive to front-to-back (FTB) motion to ON and OFF edges moving FTB and back-to-front (BTF) at 30°/s, measured using two-photon calcium imaging (data from (Salazar-Gatzimas et al., 2016)). *Right:* As at left, but for the T4 synaptic model.

The physiological properties of the inputs to T4 and T5 cells have also been characterized. At their receptive field centers, Mi1 and Tm3 cells respond quickly to visual stimuli, and provide excitatory input to T4 (Arenz et al., 2017; Behnia et al., 2014; Gruntman et al., 2018; Strother et al., 2017; Takemura et al., 2017). On the preferred direction side of the receptive field, the cells Mi4 and CT1 are ON cells with slower kinetics, likely inhibiting T4 cells (Arenz et al., 2017; Shinomiya et al., 2019; Takemura et al., 2017). On the null direction side of the receptive field, Mi9 cells are delayed OFF cells, which are likely to provide inhibitory, glutamatergic input to T4 cells (Arenz et al., 2017; Salazar-Gatzimas et al., 2018). The inputs to T5 cells similarly appear to be arranged with a fast central input and delayed flanking inputs, but whether these inputs excite or inhibit T5 is less clear (Arenz et al., 2017; Behnia et al., 2014; Shinomiya et al., 2019; Wienecke et al., 2018).

The functional properties of the T4 and T5 cells and their inputs have been interrogated using many stimulus and measurement modalities. This wealth of data has led to many different models that seek to describe the response properties of T4 and T5 cells (Arenz et al., 2017; Badwan et al., 2019; Behnia et al., 2014; Clark et al., 2011; Creamer et al., 2018; Eichner et al., 2011; Gruntman et al., 2018; Haag et al., 2016; Leong et al., 2016; Leonhardt et al., 2016; Salazar-Gatzimas et al., 2018; Salazar-Gatzimas et al., 2016; Serbe et al., 2016; Strother et al., 2017; Wienecke et al., 2018). Many measurements of T4 and T5 have demonstrated phenomenology that could not be produced by the classical HRC model. However, proposed models were most often evaluated by how they reproduced the associated dataset, rather than the full the range of phenomena in the literature. Here we ask how a minimal, constrained model reproduces T4 phenomenology (and some T5 phenomenology) from many different experiments. We compare the model to data in response to moving edges, to sinusoids, to apparent motion stimuli, to stochastic stimuli, and to natural scenes.

In this minimal model, the spatially-separated inputs to T4 are represented as three linear-nonlinear (LN) transformations of the input contrast (Dayan and Abbott, 2001). These model neurons then interact with T4 by altering the conductance of excitatory and inhibitory currents (Gruntman et al., 2018; Torre and Poggio, 1978). This construction is simple enough to allow some algorithmic intuition but incorporates greater biophysical realism than most phenomenological models. We do not fit the model to every dataset. Rather, our goal is to test the sufficiency of a minimal circuit model to account for different measured phenomena in T4 cells. This model does not contain any exotic channels or receptors, and it biophysically models the membrane voltage and intracellular calcium concentration in T4 neurons. It does not reproduce all functional properties of T4 cells, but it provides a flexible framework for understanding the sufficiency of simple circuit properties and mechanisms to describe the processing properties of T4 neurons. In cases where this model is insufficient to describe data, we suggest how model parameters might be changed to better describe the data.

## Methods

### Constructing an anatomically constrained synaptic model for T4 cells

Following proposed synaptic architectures for direction-selective computations (Gruntman et al., 2018; Torre and Poggio, 1978), we constructed an elementary motion detector based on the connectome of the *Drosophila* optic lobe. We simplified this structure to consider three inputs to a T4 cell: a delayed ND-offset OFF inhibitory input representing Mi9, a centered ON excitatory input representing Mi1 and Tm3, and a delayed PD-offset ON inhibitory input representing Mi4 (and/or CT1) (**Figure 1A**) (Strother et al., 2017; Takemura et al., 2017).

We will model these inputs to T4 cells as simple linear-nonlinear (LN) transformations of the input contrast (Behnia et al., 2014). We will further model effects of these synaptic inputs on the membrane potential of the T4 cell by changes in the conductance of excitatory and inhibitory currents (Torre and Poggio, 1978). For notational convenience, we define our model below in continuous space and time, noting as needed where adjustments are made for the discretization used in numerical simulation. We take the inputs to the model to be contrasts. We take each input to the motion detector to have an *L*_1_-normalized Gaussian spatial acceptance function

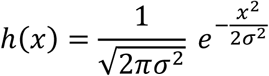

where the spatial parameter *σ* is related to the full width at half maximum (FWHM) of the acceptance function by 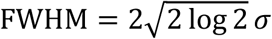. We fix FWHM = 5.7° to approximately match the spatial acceptance functions of photoreceptors in the fly eye (Stavenga, 2003). To represent the delayed inputs to the motion detector, we use the *L*_2_-normalized lowpass temporal filter

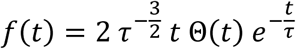

where Θ(*x*) is the Heaviside step function. To represent the non-delayed central input to the motion detector, we replace the temporal filter *f* by its derivative *ḟ*. We note that the term resulting from the distributional derivative of Θ(*t*) vanishes when *ḟ* is convolved with any signal as it is proportional to *t δ*(*t*), where *δ*(*x*) is the Dirac delta distribution. Using these filters, we define the filtered contrast signal *s* at each point in spacetime:

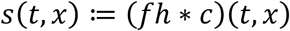

where *c*(*t*, *x*) is the input contrast and ∗ denotes spatiotemporal convolution over the appropriate domain. As taking the temporal derivative of the filtered contrast signal is equivalent to filtering with the derivative of the temporal filter, we will use the notation *ṡ* for the high-pass-filtered signal throughout. For convenient handling of spatial boundary conditions, we numerically simulate the full 360° of visual space, which is a periodic interval.

We denote the spacing between neighboring inputs as Δ. Here, we use 5° spacing so that the inputs evenly tile 360° of visual space. Then, we define the three inputs to the motion detector as rectified-linear functions of the filtered contrast signal at three points in space, mimicking the polarity-selectivity of the inputs to T4 cells:

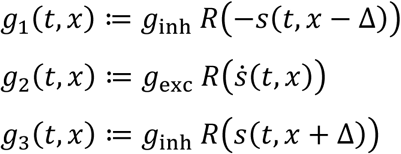

where *R*(*x*) ≔ max{0, *x*} is the ramp function and *g*_inh_ and *g*_exc_ are parameters scaling the effects of each input on the postsynaptic conductances (**Figure 1A-B**). Thus far, we have represented the conductances as linear-nonlinear (LN) transformations of the input contrast (Dayan and Abbott, 2001).

We define the membrane potential *V*_m_ of the postsynaptic cell such that the reversal potential for leak currents is 0 mV. The cell’s membrane voltage dynamics are then given as (Torre and Poggio, 1978)

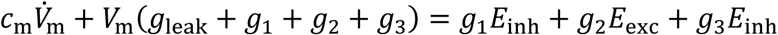

where *c*_m_ is the membrane capacitance, *g*_leak_ is the leak conductance, and *E*_inh_ and *E*_exc_ are the reversal potentials for inhibitory and excitatory currents, respectively. Neglecting capacitive currents, we solve for the pseudo-steady-state (Gruntman et al., 2018; Torre and Poggio, 1978).

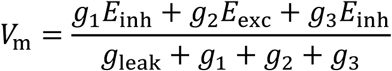

Then, we model the transformation from membrane voltage to calcium concentration *C* as a positively rectifying half-quadratic function *R*^2^(*x*) ≔ (*R*(*x*))^2^:

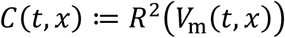

which qualitatively captures the expansive nonlinear effect of the transformation between voltage and calcium (Kato et al., 2014; Leong et al., 2016) (**Figure 1B**).

### Visual stimuli

We presented this model with spatiotemporal contrast patterns to mimic a variety of visual stimuli used in the field. Detailed mathematical descriptions of each stimulus are given in **Appendix A**. Briefly, we presented the model with moving and stationary sinusoidal gratings, with apparent motion stimuli, with stochastic stimuli including those with imposed correlations, and with natural scenes. In each case, we compared how the model responds to the published responses of T4 and T5 neurons.

### Selecting model parameters

This model uses a parameter set equal to one that was developed to explain direction-opponency in T4 cells (Badwan et al., 2019). There, we fixed the filter time constant *τ* = 150 ms to produce peak responses to PD sinusoidal gratings at ∼1 Hz (Badwan et al., 2019; Creamer et al., 2018; Maisak et al., 2013). We fix the excitatory and inhibitory reversal potentials to values of *E*_exc_ = 60 mV and *E*_inh_ = – 30 mV, which are plausible based on electrophysiological experiments (Gruntman et al., 2018). As the model membrane potential can be rewritten as

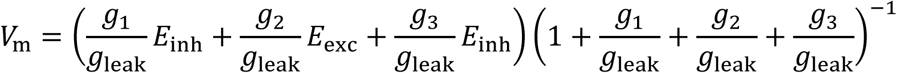

only the ratios of *g*_1_, *g*_2_, and *g*_3_ to *g*_leak_, rather than their absolute magnitudes, are relevant. We therefore express the postsynaptic conductances as non-dimensional quantities in units of *g*_leak_, leaving *g*_exc_/*g*_leak_ and *g*_inh_/*g*_leak_ as the model’s two free parameters. The procedure used to select the values of these parameters is described in detail in **Appendix B**. As shown previously (Badwan et al., 2019), there exists a broad region of parameter space for which this model displays responses to sinusoid gratings with a temporal frequency of 1 Hz and a spatial wavelength of 45° consistent with those measured in T4 and T5 cells. We note that our choice of filter normalization, which differs from that in the previous use of this model (Badwan et al., 2019), affects the parameter values chosen, as it scales *g*_1_, *g*_2_, and *g*_3_ relative to *g*_leak_. **Table 1** summarizes the model parameter values used in all simulations.

**Table 1:** Parameter values used in all simulations.

| Parameter | Value |
| --- | --- |
| Photoreceptor spacing $\Delta$ | $5^\circ$ |
| Photoreceptor spatial FWHM | $5.7^\circ$ |
| Temporal filter time constant $\tau$ | 0.150 s |
| Spatial sampling interval $\Delta x$ | $0.5^\circ$ |
| Temporal sampling interval $\Delta t$ | 1/240 s |
| Leak current reversal potential $E_{\text{leak}}$ | 0 mV |
| Excitatory current reversal potential $E_{\text{exc}}$ | +60 mV |
| Inhibitory current reversal potential $E_{\text{inh}}$ | -30 mV |
| Excitatory to leak conductance ratio $g_{\text{exc}}/g_{\text{leak}}$ | 0.1 |
| Inhibitory to leak conductance ratio $g_{\text{inh}}/g_{\text{leak}}$ | 0.3 |

### In vivo two-photon calcium imaging in T4 cells

Most of our comparisons relate the synaptic model’s responses to published data, but we also compare the model to a new dataset of T4 cell responses to glider stimuli. The protocol for two-photon calcium imaging in T4 cells matches published methods (Badwan et al., 2019) and used Psychtoolbox (Brainard, 1997; Kleiner et al., 2007; Pelli, 1997) to present stimuli on a panoramic visual display (Creamer et al., 2019). The glider stimuli presented during these measurements are described in **Appendix A**. Net responses were computed as the difference in responses to stimuli moving in the preferred and null directions of each T4 region of interest, and then averaged within each fly. Non-parametric two-sided Wilcoxon signed-rank tests were used to test whether median net responses differed significantly from zero (Hollander et al., 2013). For statistical purposes, each individual fly was considered to be an independent sample.

### Numerical methods

Numerical simulations were conducted using Matlab 9.6 (R2019a) (The MathWorks, Natick, MA, USA). For stimuli containing randomly-generated components, responses were averaged over 1000 realizations, and bootstrapped 95% confidence intervals for the mean were computed using the bias-corrected and accelerated percentile method (Efron, 1987).

## Results

### The synaptic model reduces to HRC-like terms

To gain intuition about the operation of the T4 synaptic model, we consider its expansion in the small-input limit. To do so, we approximate the ramp function nonlinearity with a smooth function that represents a soft rectifier, which can be approximated by a linear function for small inputs (Fitzgerald and Clark, 2015). In particular, letting *s*_1_(*t*) ≔ *s*(*t*, *x* − Δ), *s*_2_(*t*) ≔ *s*(*t*, *x*), and *s*_3_(*t*) ≔ *s*(*t*, *x* + Δ), and defining the non-negative constants *α* ≔ |*g*_inh_*E*_inh_/*g*_leak_| and *γ* ≔ |*g*_exc_*E*_exc_/*g*_leak_|, we have, to lowest order in the inputs,

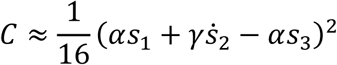

which may be rewritten as

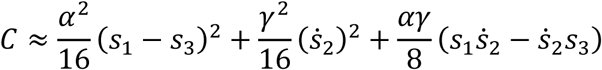

This expansion represents a motion-energy approximation of the model. The first term in this expansion is a finite-difference approximation to a spatial derivative, while the second term is a temporal derivative at the center of the model’s receptive field. The third term, which is the only direction-selective term, is the subtraction of two offset correlators with opposite directional tuning. This subtraction step provides some intuition for why this model mimics some properties of a fully-opponent HRC model (Badwan et al., 2019). This same direction-selective term also appears in the second-order expansion of the membrane voltage. Because this expansion of the model is only to second-order, it is invariant under contrast inversions, and cannot account for properties like ON-edge selectivity (Clark et al., 2014; Fitzgerald and Clark, 2015; Fitzgerald et al., 2011). Though this simple description does not capture all properties of the synaptic model, it provides intuition for the sensitivity of the model to certain stimulus features.

### The synaptic model is strongly ON/OFF-edge- and direction-selective

T4 and T5 neurons are distinguished by the fact that T4 cells respond to ON-edges while T5 cells respond to OFF-edges (Maisak et al., 2013). We first compared the ON/OFF edge- and direction-selectivity of our T4 synaptic model to responses measured using two-photon calcium imaging in T4 cells sensitive to front-to-back (FTB) motion. Like T4 FTB cells, our synaptic model responded strongly to an ON edge moving in the FTB direction, but displayed little or no response to OFF edges moving in the FTB direction or to edges of either polarity moving in the back-to-front (BTF) direction (**Figure 1C**) (Maisak et al., 2013; Salazar-Gatzimas et al., 2016).

### The spatiotemporal tuning of the synaptic model is consistent with that of T4 cells

Sinusoid grating stimuli are a common tool for characterizing direction-selective computations. Responses to these stimuli have been used to suggest that the membrane voltage of T5 cells is a nearly linear transformation of the visual input (**Figure 2A**) (Wienecke et al., 2018). In the synaptic model, the membrane voltage is a nonlinear function of the input contrast because the inputs are first rectified and then interact nonlinearly. We applied the same linearity testing protocol to our model membrane voltage, constructing predictions for responses to PD and ND drifting gratings from the responses to counterphase gratings (see **Appendix A**) (Jagadeesh et al., 1993; Wienecke et al., 2018). The responses of the T4 synaptic model to drifting gratings were similar to those predicted by a linear model for membrane voltage (**Figure 2A**). Thus, even a nonlinear system like the T4 synaptic model may appear reasonably linear by this protocol.

**Figure 2:**
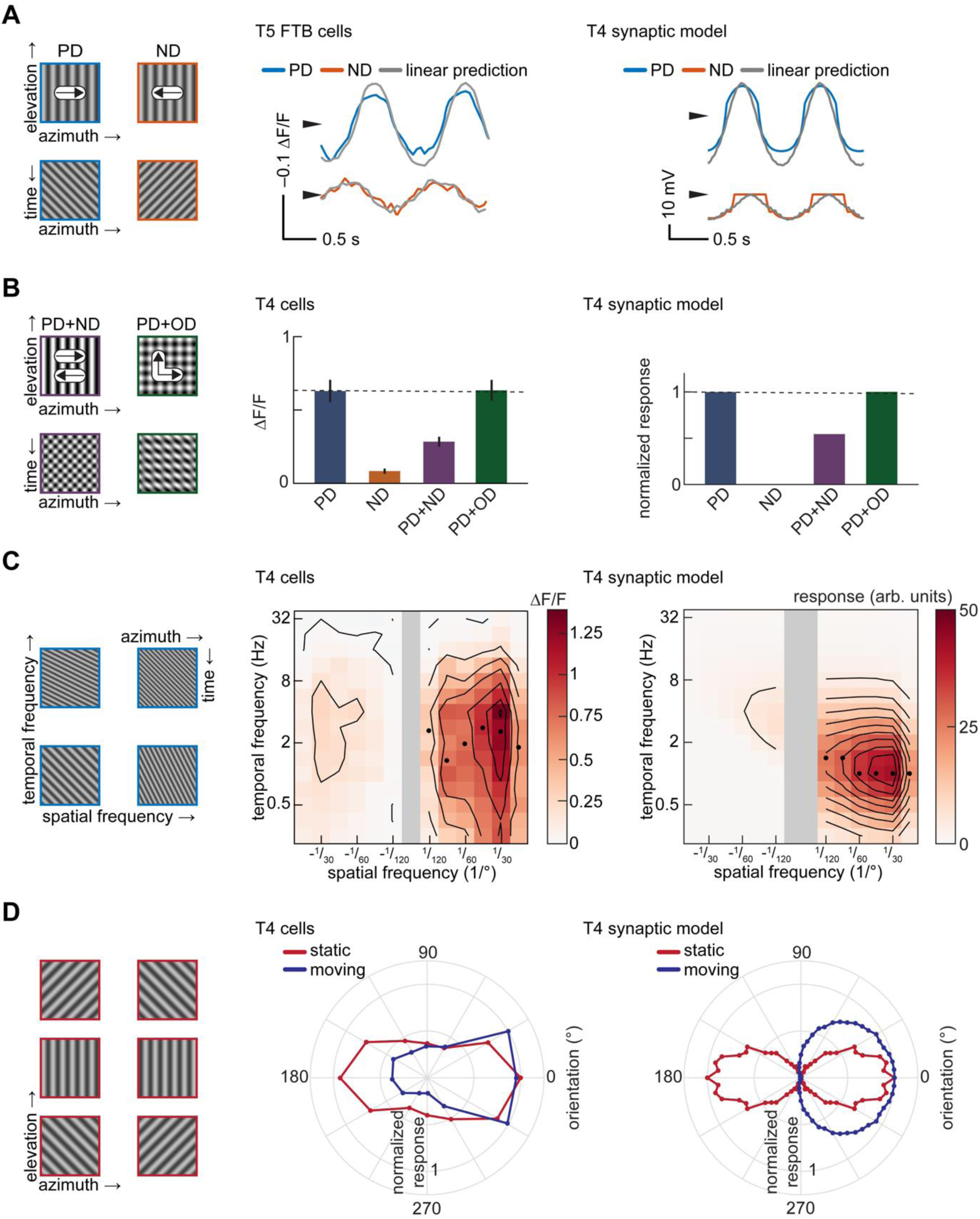
The T4 synaptic model reproduces sinusoidal grating responses measured in T4 cells. A. *Left:* Images and kymographs of sinusoid gratings drifting in the preferred (PD) and null (ND) directions. *Center:* Membrane voltage of T5 FTB cells to 1 Hz, 25° drifting gratings compared with linear predictions from contrast-modulated counterphase gratings, measured using voltage indicators (data from (Wienecke et al., 2018)). *Right:* As at center, but for voltage responses of the T4 synaptic model. The model has coefficients of determination for PD and ND of 0.92 and 0.82. B. Mean responses to 1 Hz, 45° sinusoid gratings. *Left:* Images and kymographs of composite sinusoid gratings containing PD and ND motion or PD and orthogonal direction (OD) motion. *Center:* Mean responses of T4 cells to drifting gratings, measured using a calcium indicator (data from (Badwan et al., 2019)). Error bars indicate ±1 SEM. *Right:* As at center, but for calcium responses of the T4 synaptic model. C. Spatiotemporal frequency tuning. *Left:* Kymographs of sinusoid gratings with varying spatiotemporal frequency content. *Center:* Spatiotemporal frequency tuning of T4 cells (data from (Creamer et al., 2018)). Black circles indicate the temporal frequency at which the maximum response at a given spatial frequency is attained. *Right:* As at center, but for the T4 synaptic model. D. Orientation and direction tuning. *Left:* Images of oriented sinusoid gratings. *Center:* Orientation tuning of T4 and T5 cells with static gratings (data from (Fisher et al., 2015)) and direction tuning of T4 cells with drifting gratings (data from (Maisak et al., 2013)). The orientation of a static grating is defined by the vector normal to the apparent edges, the same definition as for moving gratings (see **Appendix A**). *Right:* As at center, but for the T4 synaptic model.

T4 and T5 cells display direction-opponent average calcium responses to sinusoid gratings (**Figure 2B**) (Badwan et al., 2019). This property means that the average response to PD motion is reduced by the addition of ND motion, imposing a strong constraint on models for the direction-selective computation. In particular, it implies that linear-nonlinear models with expansive nonlinearities cannot account for the response properties of these cells (Badwan et al., 2019). A variant of this synaptic model was proposed to account for these direction-opponent responses (Badwan et al., 2019). This model reproduces the strong suppression when ND motion is added to PD motion without substantial enhancement when orthogonal-direction (OD) motion is added to PD motion (**Figure 2B**).

T4 and T5 cells are tuned to the temporal frequency of sinusoidal stimuli (**Figure 2C**) (Creamer et al., 2018). This means that the mean neural response is maximal at a single temporal frequency, independent of the wavelength. This property also applies to measurements of fly behavior (Creamer et al., 2018; Kunze, 1961) and is consistent with the classical, fully-opponent HRC. We presented the T4 synaptic model with drifting gratings of different spatial and temporal frequencies to find the mean response to each. The model response was strongly temporal-frequency-tuned (**Figure 2C**). To quantify the temporal-frequency-tuning, we asked how much of the variance in this surface was accounted for by the product of one function of temporal frequency and one fuction of spatial frequency response (Creamer et al., 2018; Priebe et al., 2006). Such a separable model accounted for 99% of the variance in the response (see **Appendix A**, **Figure 2C**). Because of our choice of parameters, the input temporal filters in this model produce peak responses at around 1 Hz, lower than the roughly 2-4 Hz peak measured in these T4 cells.

T4 and T5 cells respond to static gratings with amplitudes that depend on the grating orientation (Fisher et al., 2015) (**Figure 2D**). The preferred orientation (defined by the vector normal to the edges in a static grating) approximately matches the preferred direction of motion of these cells (Maisak et al., 2013). The convention we use here for defining the orientation of a static grating is rotated 90° relative to that used in the original study, which defined orientation in terms of vectors parallel, rather than normal, to the edges (Fisher et al., 2015) (see **Appendix A**). When the T4 synaptic model was presented with both static and drifting gratings of many different orientations, it reproduced the orientation tuning observed experimentally for both static and moving gratings (**Figure 2D**). The model was more selective for both orientation and direction than the T4 cell measurements.

### The synaptic model reproduces the selectivity of apparent motion responses in T4 cells

In addition to sinusoid gratings, apparent motion stimuli are a useful tool for investigating direction-selective systems. These stimuli decompose visual motion into summations of simpler spatiotemporal patterns, which can provide strong intuition into the motion computation (Barlow and Levick, 1965).

Electrophysiological measurements of T4 cells have shown fast depolarization and delayed, offset hyperpolarization in response to a small flashed white bar placed on a gray background (Gruntman et al., 2018) (**Figure 3A**). The synaptic T4 model displayed qualitatively consistent responses to the same stimulus (**Figure 3A**). The positive lobe in the model is narrower than in the electrophysiological recording; this is likely because the true central input to T4 has a wider receptive field than in our model (Behnia et al., 2014; Takemura et al., 2013). Consistent with electrophysiology, the OFF input to the T4 model is not visible under this analysis because it was rectified with a threshold at mean gray (zero contrast).

**Figure 3:**
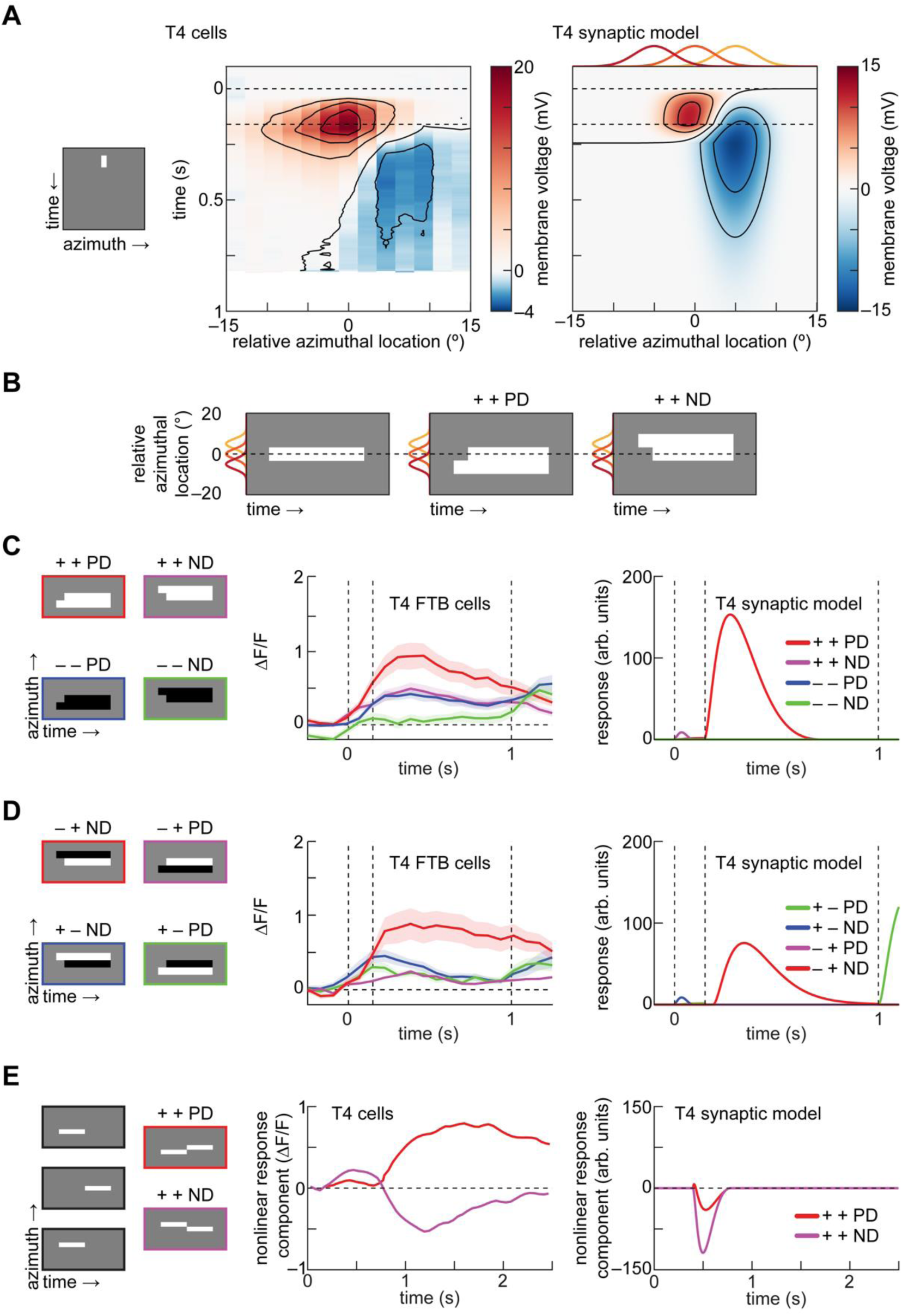
The T4 synaptic model reproduces the spatial organization and selectivity of apparent motion responses in T4 cells. A. Responses to a single white bar flashed at different spatial locations. *Left:* Kymograph of 2° white bar presented for 160 ms. *Center:* Membrane voltage of T4 cells to flashed white bars, measured using electrophysiology (data from (Gruntman et al., 2018)). *Right:* As at center, but for the T4 synaptic model. Red, orange, and yellow lines indicate the spatial acceptance functions of the three model inputs. B. Apparent motion stimuli are aligned such that the lagging bar is located at the center of the receptive field (Salazar-Gatzimas et al., 2018). The leading bar is presented at time zero and lasts for 1 second, and the lagging bar is presented 150 ms later. Each bar subtends 5° of visual angle. Red, orange, and yellow lines indicate the spatial acceptance functions of the three inputs. C. Responses to phi apparent motion stimuli, aligned as in (B). *Left:* Kymographs of all four possible phi apparent motion stimuli. *Center:* Responses of T4 FTB cells to all four phi apparent motion stimuli, measured using two-photon calcium imaging (data from (Salazar-Gatzimas et al., 2018)). Error patches indicate ±1 SEM. *Right:* As at center, but for the T4 synaptic model. D. As in (C), but for reverse-phi apparent motion stimuli, in which the sequentially presented bars have opposite contrasts. E. Assessing PD enhancement and ND suppression. *Left:* Kymographs of linear decomposition of flashed apparent motion stimuli, with 4.5°-wide white bars presented sequentially for 400 ms each. *Center:* Nonlinear response component, defined as the residual of the linear prediction, measured using a calcium indicator (data from (Haag et al., 2016)). *Right:* As at center, but for the T4 synaptic model.

Since the synaptic model reproduced T4 cell voltage responses to flashed bars, we sought to characterize its responses to apparent motion stimuli composed of pairs of bars offset in spacetime (Salazar-Gatzimas et al., 2018). These stimuli can induce in humans and in flies the “reverse-phi” motion illusion, in which a reversal of contrast polarity induces a motion percept in the direction opposite the stimulus displacement (Anstis, 1970; Clark et al., 2011; Hassenstein and Reichardt, 1956). We aligned these stimuli so that the temporally-delayed bar is placed at the center of the receptive field (**Figure 3B**) (Salazar-Gatzimas et al., 2018). T4 cells respond maximally to one phi and one reverse-phi apparent motion stimulus out of eight possible pairings (Salazar-Gatzimas et al., 2018). The synaptic model reproduced this selectivity (**Figure 3C-D**).

Various groups have assessed nonlinear enhancement or suppression of PD and ND apparent motion stimuli relative to linear decompositions. This analysis can be misleading because it does not allow one to uniquely characterize the nonlinearity as ‘enhancing’ or ‘suppressing’, since there exist an infinite number of linear decompositions of a given stimulus (Salazar-Gatzimas et al., 2018). Despite this difficulty, such analyses have been applied as an intuitive way to try to understand direction-selective computations (Barlow and Levick, 1965; Fisher et al., 2015; Gruntman et al., 2018; Haag et al., 2016).

In T4 cells, an analysis of responses to sequential bars has indicated that calcium signals include both PD enhancement and ND suppression relative to a linear prediction from the responses to individual bars (Haag et al., 2016) (**Figure 3E**). Our model failed to reproduce this result, showing only suppression of ND motion under this analysis (**Figure 3E**). This discrepancy could be influenced by the timescale of this stimulus, which is far longer than the 150 ms offset used in the apparent motion stimuli in (**Figure 3C-D**). Additionally, previous theoretical work has shown that disinhibition can generate PD enhancement in similar models (Borst, 2018; Torre and Poggio, 1978); the choice of thresholds in this model did not permit flanking disinhibition with ON stimuli.

### The synaptic model does not reproduce the fast timescale tuning of T4 cells

A third approach to characterizing direction-selective signals has been to apply stochastic stimuli with specified correlation structure. Responses to uncorrelated stimuli can be used to generate an unbiased estimate of a system’s linear receptive field (Chichilnisky, 2001). By using reverse-correlation and uncorrelated stimuli to extract spatiotemporal receptive fields, T4 cells have been characterized by oriented linear receptive fields with a central excitatory lobe and a delayed, offset inhibitory lobe (**Figure 4A**) (Leong et al., 2016; Salazar-Gatzimas et al., 2016). The T4 synaptic model generates the same shape of receptive field (**Figure 4A**). However, in the model, the inhibitory lobe lasts longer than that measured in T4 cells, and the tuning of the model was slower overall.

**Figure 4:**
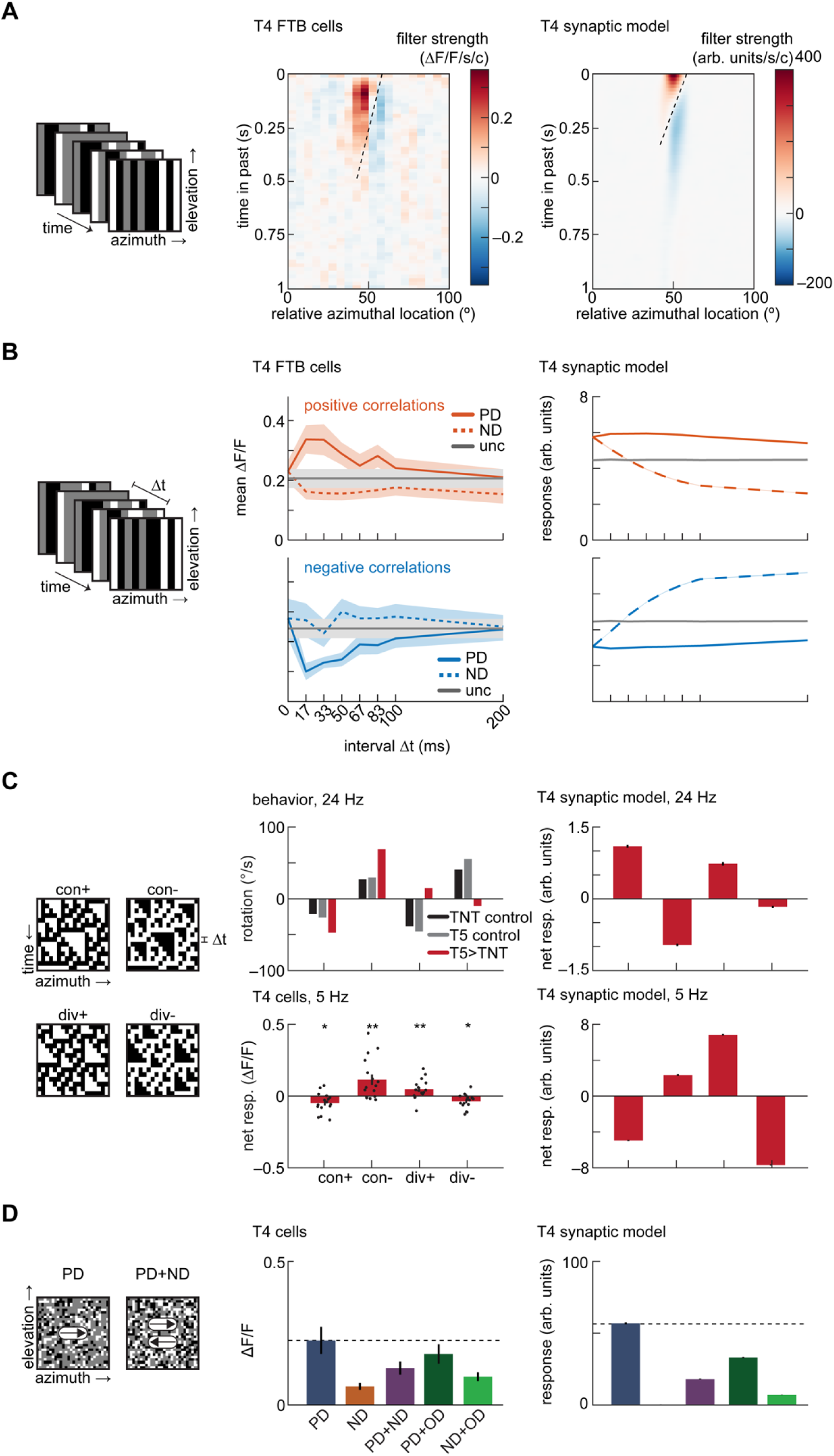
The minimal T4 synaptic model is not sufficient to reproduce the fast-timescale tuning of T4 cells. A. Linear receptive field measurements. *Left:* Schematic depiction of binary, uncorrelated spatiotemporal noise. *Center:* Linear receptive field of T4 FTB cells (data from (Salazar-Gatzimas et al., 2016)). *Right:* As at center, but for the T4 synaptic model. B. Correlation interval receptive field measurements. *Left:* Schematic depiction of ternary noise containing pairwise correlations at a specified interval Δ*t*. *Center:* Responses of T4 FTB cells to positive and negative correlations, measured using a calcium indicator (data from (Salazar-Gatzimas et al., 2016)). Error bars indicate ±1 SEM. *Right:* As at center, but for the T4 synaptic model. Error patches, which are barely visible, indicate 95% confidence intervals of the mean, which is variable due to the stochastic stimulus. C. Triplet correlation sensitivity. *Left:* Kymographs of three-point glider stimuli containing positive and negative triplet correlations. *Center top:* Turning behavioral responses to three-point gliders updated at 24 Hz of flies with the synaptic outputs of T5 cells silenced (data from (Leonhardt et al., 2016)). Positive rotations correspond to the direction of the displacement of the spatial mean location of each triplet. *Center bottom:* Net responses of T4 cells to three-point gliders updated at 5 Hz, measured using a calcium indicator (see **Methods**). Asterisks indicate that median net response differs from zero at the p < 0.05 (*) or p < 0.01 (**) level by a Wilcoxon signed-rank test with N = 16 flies. Exact p-values are p = 0.0174, 0.0061, 0.0097, and 0.0131 for con+, con-, div+, and div-, respectively. Error bars indicate ±1 SEM over flies, and black circles indicate individual per-fly means. *Right:* As at center, but for the T4 synaptic model. Error bars indicate 95% confidence intervals of the mean. D. Responses to rigidly translating stimuli with stochastic checkerboard patterns. *Left:* Images of random checkerboard stimuli. *Center:* Mean responses of T4 cells to checkerboard stimuli translating at 100°/s, measured using a calcium indicator (data from (Badwan et al., 2019)). Error bars indicate ±1 SEM. *Right:* As at center, but for the T4 synaptic model. Error bars indicate 95% confidence intervals of the mean.

Responses to stochastic stimuli containing precise pairwise spatiotemporal correlations have revealed fast-timescale tuning in T4 cells (Salazar-Gatzimas et al., 2016). In measurements of T4 and T5, the cells could discriminate between spatiotemporal correlations with delays of 0 and 15 ms (**Figure 4B**). We presented the synaptic model with stimuli containing pairwise spatiotemporal correlations at different temporal delays. The model was direction-selective and responded to both positive and negative correlations, as in the cellular measurements. However, the model did not reproduce the fast timescale discrimination between delays (**Figure 4B**). Furthermore, the synaptic model showed strong suppression of ND-oriented positive correlations and enhancement of ND-oriented negative correlations, which was not observed in the data.

Behaving *Drosophila* respond direction-selectively to correlations higher than second-order (Clark et al., 2014; Leonhardt et al., 2016). This cannot be explained by models that compute pairwise correlations in the stimulus, such as the HRC and motion energy model. The sensitivity to higher-order correlations has been assessed using three-point glider stimuli, which contain precise third-order correlations (Hu and Victor, 2010) (**Figure 4C**). The net responses of T4 cells to these stimuli have previously been inferred from behavioral measurements in *Drosophila* with the synaptic outputs of T5 cells silenced, using gliders updated at 24 Hz (Leonhardt et al., 2016). We used *in vivo* two-photon calcium imaging to measure directly the responses of T4 cells to three-point gliders updated at 5 Hz, and found that the signs of the net responses were consistent with those measured in behavior with T5 cells silenced (**Figure 4C**, see **Methods** and **Appendix A** for details).

With an update rate of 24 Hz, the synaptic model correctly predicted the signs of net responses to diverging gliders measured in imaging and behavior, but predicted the wrong converging glider responses (**Figure 4C**). At 5 Hz, the synaptic model correctly predicted the signs of both converging and diverging glider responses, but not the relative magnitudes. Thus, the glider responses in T4 appear relatively insensitive to the glider timescale (24 vs. 5 Hz), but the model’s response depends strongly on the input timescales.

T4 and T5 cells have been shown to display strongly direction-selective responses to rigidly-translating stimuli consisting of black and white squares placed at random on a gray background (see **Appendix A**) (Badwan et al., 2019). When two such stimuli that move in opposite directions are superimposed, they generate transparent motion percepts in primates (Qian et al., 1994), and they generate responses in T4 that are reduced compared to presenting PD stimuli alone (Badwan et al., 2019). The synaptic model qualitatively reproduced these responses (**Figure 4D**). In particular, the responses of T4 cells are suppressed more strongly under the addition of ND motion than under the addition of OD motion, a feature that is reproduced by the synaptic T4 model (**Figure 4D**). Therefore, as in T4 cells, the selective direction-opponency observed in the model persists even with stimuli containing multiple spatiotemporal frequencies.

### The T4 synaptic model provides decorrelated channels for naturalistic motion

Beyond artificial stimuli, natural scenes have been used to investigate the performance of direction-selective signals generated by models, behavior, and neurons (Badwan et al., 2019; Chen et al., 2019; Dror et al., 2001; Fitzgerald and Clark, 2015; Leonhardt et al., 2016; Salazar-Gatzimas et al., 2018; Straw et al., 2008). We therefore sought to investigate the performance of the T4 synaptic model in natural motion processing. To do so, we presented it with rigidly-translating scenes from a database of natural images (see **Appendix A, Figure 5A**) (Meyer et al., 2014) (Badwan et al., 2019; Chen et al., 2019; Fitzgerald and Clark, 2015; Meyer et al., 2014; Salazar-Gatzimas et al., 2018). Though the structure and properties of inputs to T5 cells are known to differ from the inputs to T4 (Serbe et al., 2016; Shinomiya et al., 2019), to make an OFF-edge selective channel we created a ‘T5’ model by simply inverting the ON/OFF selectivity of the inputs to our T4 synaptic model. This is intended merely to be an OFF-selective channel for the purposes of comparing T4 and potential T5 cell responses. The resulting four channels displayed strongly direction-selective average responses to translating natural scenes (**Figure 5B**).

**Figure 5:**
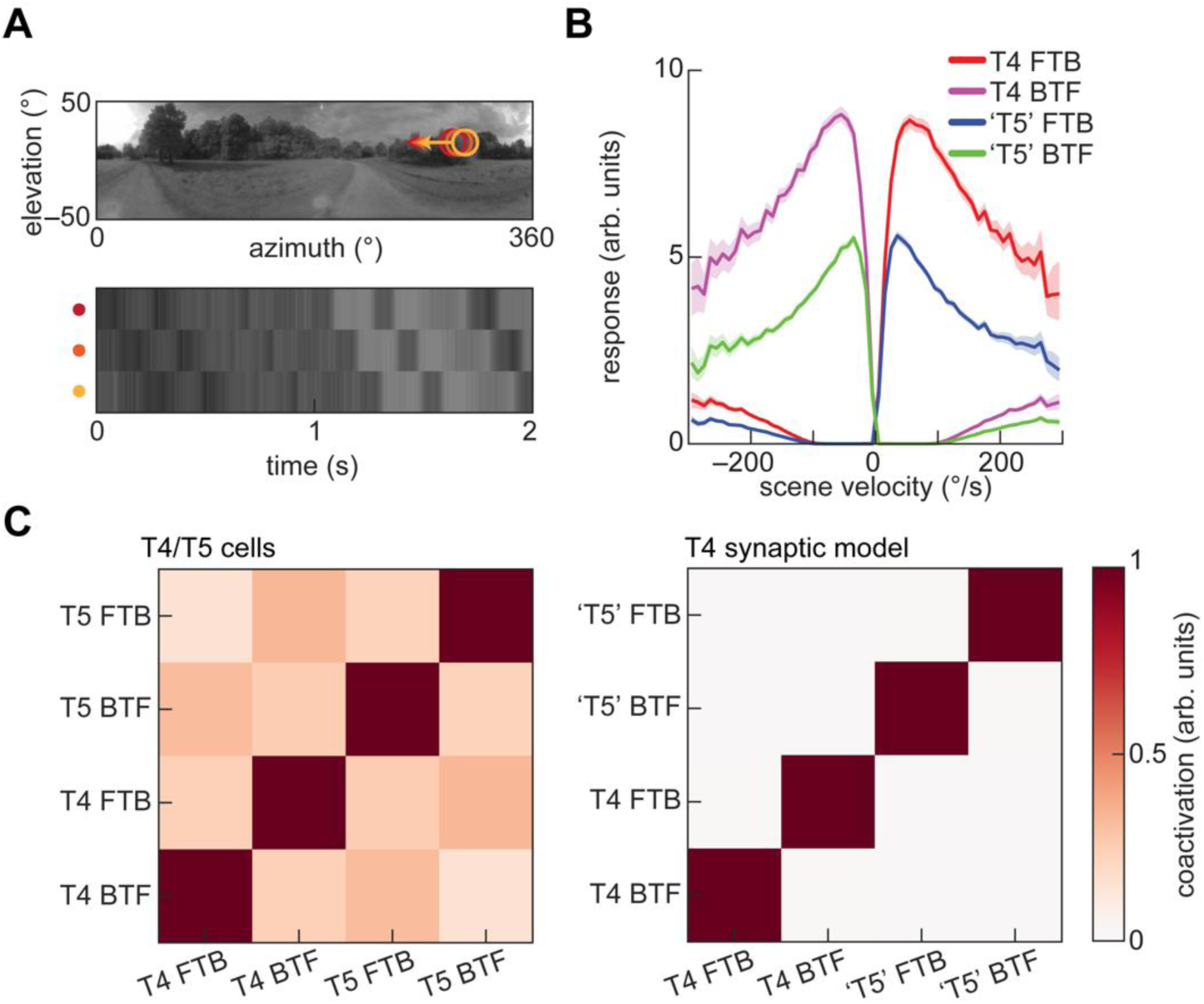
The T4 synaptic model generates decorrelated signals in response to naturalistic motion. A. A rigidly rotating panoramic natural scene, with three spatially offset input signals as a function of time. B. Average responses of the T4 synaptic model and a ‘T5’ variant to naturalistic motion constructed by rigidly translating natural scenes at a variety of velocities. Error patches indicate 95% confidence intervals of the mean. ‘T5’ cells were constructed by sign-inverting the inputs to the minimal T4 synaptic model. C. Decorrelation of channels with naturalistic motion. *Left:* Coactivation matrix of T4 and T5 cells in response to rigidly translating natural scenes (data from (Salazar-Gatzimas et al., 2018)). *Right:* As at left, but for the T4 synaptic model and ‘T5’ variant. Coactivation was computed for an ensemble of (image, velocity) pairs, in which the velocity was chosen from a Gaussian distribution with zero mean and 100°/s standard deviation.

Measured responses of T4 and T5 cells to translating natural scenes are decorrelated, so that only one channel is active at once (Salazar-Gatzimas et al., 2018). The synaptic models of T4 and ‘T5’ also generated highly decorrelated responses, with the coactivation matrix of the four channels being nearly diagonal (**Figure 5C**). Such decorrelated parallel channels may provide a convenient representation of motion signals (Salazar-Gatzimas et al., 2018).

## Discussion

An anatomically constrained synaptic model suffices to reproduce many, but not all, of the properties of *Drosophila* T4 cells. This model reproduces the direction-opponency, temporal-frequency-tuning, orientation-tuning, and phi/reverse-phi selectivity measured in T4 cells (**Figures 2–4**). When applied to a naturalistic velocity estimation task, it produces decorrelated signals similar to those measured in T4 and T5 neurons (**Figure 5**). However, it fails to reproduce the PD enhancement and fast-timescale tuning observed in T4 cells (**Figures 3–4**). Moreover, though it is sensitive to triplet correlations in its input, it fails to reproduce them on the same timescales as observed in the data (**Figure 4**). In short, this simple synaptic model is sufficient to reproduce several distinct properties of T4 cells, but cannot account for several observations.

### Minimal models and levels of understanding

Here, we asked whether a minimal synaptic model could qualitatively reproduce features of T4 cell responses. The minimal model required no exotic neurotransmitter receptors or interactions, and was based on simple synaptic conductances. The simplifications sufficient to explain different phenomena will depend strongly on the features one seeks to reproduce, and on the desired level of fidelity. However, minimal models are useful precisely because they can be relatively straightforward to analyze.

Marr famously proposed different levels of understanding neural circuitry, including an algorithmic level and a mechanistic level (Marr and Poggio, 1976). As we drive towards a deeper understanding of the visual motion circuit in the fly, the levels of algorithm and mechanism can appear increasingly blurred. It is hard to define what distinguishes the mechanistic circuit description presented here from a detailed algorithm-level description of the computation. However, it remains important to connect proposed mechanistic models to high-level descriptions of the system such as the HRC. This is because the high-level descriptions of computations provide a level of intuition for the behavior of the system that a more intricate model cannot. Moreover, the HRC explains a wide variety of neural and behavioral data in flies (Borst and Egelhaaf, 1989; Yang and Clandinin, 2018), so an HRC-like algorithm must be a limiting case of any proposed mechanistic model (Potters and Bialek, 1994).

### Sufficiency of models

Many details of the function of the early visual system were neglected in this model. For instance, the filter shapes in neurons leading into the model T4 cell have been well-characterized (Arenz et al., 2017; Behnia et al., 2014), but this model used simple exponential filters. Lateral inhibition is widely documented in the early fly visual system (Arenz et al., 2017; Freifeld et al., 2013; Meier et al., 2014), but this model used simple Gaussian spatial acceptance functions without lateral inhibition. The synapses that feed into the medulla neurons that synapse onto T4 are likely to have complex, nonlinear processing properties (Yang et al., 2016), yet we modelled the entire input pathway as a purely linear filter. The rectifications of neural responses upstream of T4 are imperfect (Behnia et al., 2014; Salazar-Gatzimas et al., 2018), but this model used simple threshold-linear rectifiers. The fly eye possesses neurons that feedback onto earlier stages and create reciprocal interactions between neurons (Takemura et al., 2013; Takemura et al., 2017), but this model is entirely feedforward.

Despite these approximations, the synaptic T4 model presented here is sufficient to qualitatively match a variety of T4 neuron responses. Adding some of these neglected details into a model may make it sufficient to reproduce other features of T4 responses. This provides a method for understanding which details of processing are related to which response features in T4 cells: one may ask how different details of the system affect the sufficiency of a model to reproduce specific downstream response properties. As the field acquires more and more detailed information about the motion detection circuitry, this sort of analysis will be critical to understand the functional role of different properties.

One might naturally ask whether the synaptic model presented here might be further simplified without sacrificing its ability to account for the response properties of T4 cells. As described in **Appendix C**, a simplified linear-nonlinear cascade (LNLN) representing the numerator of the biophysical nonlinearity can generate some, but not all, of the properties of the full model.

### Flexibility in extending this minimal synaptic model

In selecting parameter values for this synaptic model, we sought to reproduce only a few properties of T4 cells: a temporal frequency maximum of 1 Hz and a direction-opponent average responses to sinusoid gratings with a temporal frequency of 1 Hz and a spatial wavelength of 45° (**Appendix B**) (Badwan et al., 2019). To capture a larger subset of the measured properties of T4 cells, one could optimize the parameters of the model capture many response properties (Deb, 2014). Such a solution would provide information about the maximal ability of this synaptic model to reproduce the properties of T4 cells, but it seems unlikely to provide insight into the predictive power of the core features of the model.

The organization of this model allows for several clear tuning mechanisms. First, the temporal filters could be modified to better match measured filters (**Figure 2**). Second, the degree to which inhibition is shunting or hyperpolarizing can be adjusted by changing the reversal potential of inhibitory currents. This could effectively hide inhibition under some stimuli and measurements. Third, it is clear that to better represent preferred direction enhancement, the threshold for the OFF-inhibitory input could be changed (**Figure 3**) (Borst, 2018). This would allow disinhibition of Mi4 to change the gain for the central input.

In the model analyzed here, we chose all thresholds of the input LN models to be zero. This effectively ignores contrast asymmetries in the natural world (Geisler, 2008), which have been used to understand many functional properties of motion detectors in flies (Chen et al., 2019; Clark et al., 2014; Fitzgerald and Clark, 2015; Fitzgerald et al., 2011; Leonhardt et al., 2016; Salazar-Gatzimas et al., 2018). Changing these thresholds to optimize for natural scene motion estimation might also generate a parameter set that better captures responses to triplet correlations (**Figure 4**) (Fitzgerald and Clark, 2015). In short, the synaptic model presented here is highly flexible and extensible, and uses only simple, known biophysical mechanisms.

For the sake of simplicity, we have used single delay and non-delay lines in this work (**Figure 1A**). However, T4 cells receive fast excitatory input at the center of their receptive fields from both Mi1 and Tm3 cells, and delayed OFF inhibitory input offset in the PD from both Mi4 and CT1 cells (Shinomiya et al., 2019; Takemura et al., 2017). Dissecting how information from these parallel channels is used, particularly if it is nonlinearly combined, will be important in developing a full understanding of the direction-selective computation performed by T4 cells.

### Modelling temporal processing

The model presented here failed to capture some of the fast-timescale tuning measured in T4, including in its responses to pairwise and triplet spatiotemporal correlations (**Figure 4**). In this minimal model, we represented all temporal processing by linear filters. However, the temporal processing upstream of T4 cells involves nonlinear and adaptive mechanisms, which can affect temporal response properties (Howard et al., 1984; Zheng et al., 2006). Thus far, the study of nonlinear mechanisms in the fly visual system has focused on static nonlinear effects such as rectification (Behnia et al., 2014; Yang et al., 2016) and on nonlinear interactions between linearly filtered signals (Borst et al., 2005; Fitzgerald and Clark, 2015). The inclusion of nonlinear effects on the dynamics themselves may be necessary to accurately capture the temporal processing upstream of T4 cells. As a first step towards experimentally understanding adaptation in this circuit, one might characterize the temporal kernels of the inputs to T4 cells with high resolution (Mano et al., 2019; Yang et al., 2016) and study how their properties depend on stimulus statistics and history (Baccus and Meister, 2002; Kim and Rieke, 2001; Rieke, 2001). Only a few models have focused on these sorts of changes in processing dynamics (Clark et al., 2013). Though the analysis of dynamic temporal nonlinearities is complex, incorporating them into models may provide insight into how fast timescale tuning of T4 cells arises.

### A T5 synaptic model

In this work, we used a sign-inverted version of our T4 synaptic model to represent the OFF-edge-selective T5 cells. This representation would correspond to a first-order direction-selective cell that receives OFF excitatory input at the center of its receptive field, delayed OFF inhibitory input offset in its preferred direction, and delayed ON input offset in the null direction. Such a model would correctly predict the selective responses of T5 cells to phi and reverse-phi apparent motion stimuli (Salazar-Gatzimas et al., 2018). However, the functional and anatomical structure of the inputs to T5 cells suggests that it receives only OFF inputs (Serbe et al., 2016; Shinomiya et al., 2019). Somehow, however, signals in T5 cells are sensitive to both contrast increments and decrements (Salazar-Gatzimas et al., 2018; Wienecke et al., 2018). Further study of the physical and functional connectome of the OFF-edge motion pathway, will be required to elucidate how the direction-selective computations in T4 and T5 cells differ.

### Relationships to mammalian visual systems

The organization of the fly’s visual motion detection circuits bear striking similarities to those in mammalian retina in their anatomy, circuitry, and algorithmic processing (Borst and Helmstaedter, 2015; Clark and Demb, 2016; Sanes and Zipursky, 2010). In mammalian retina, the earliest direction-selective signals are generated by starburst amacrine cells (SACs), which are also tuned to ON- and OFF-edges (Euler et al., 2002; Famiglietti Jr, 1983). It appears that SACs may receive inputs that are differentially delayed (Fransen and Borghuis, 2017; Kim et al., 2014), similar to the inputs to T4 cells. It would be interesting to investigate how much SAC phenomenology that mechanism alone could account for, when linked to simple biophysical mechanisms. As in this study, it could provide insight into where the circuit understanding is lacking, especially when complex stimuli are used to probe SAC function (Chen et al., 2016).

It is notable that the ON-ON-OFF spatial organization of T4 inputs (**Fig. 1A**) is almost identical to a model proposed to explain cortical responses to pairwise correlations (Mo and Koch, 2003). This suggests there may be deep parallels between T4 and T5 and cortical motion processing steps. Models for fly and cortical direction-selectivity have traditionally differed in whether they assume discrete inputs (fly, HRC-like models) or more continuous inputs (cortex, motion-energy-like spatiotemporal filtering). If synaptic interactions are considered, then continuous linear filters cannot be applied, and models must incorporate the discrete receptive fields of the inputs to a cell. It would be interesting to ask how such conductance models fare in predicting cortical responses; the statistical nature of cortical connections make it more difficult to make a general model of this type.

In this synaptic model of T4 cell function, we have paired known connectivity with measured physiology and simple biophysics to predict many circuit processing properties. This allows us to define where such a model succeeds and where it fails. This represents progress towards the ultimate goal of understanding this circuit at all levels, from utility to algorithm to mechanism.

## Author contributions

JAZV and DAC conceived of numerical experiments. JAZV performed numerical simulations and analyzed the model. BAB acquired calcium imaging data. JAZV and DAC wrote the paper.

## Acknowledgements

We thank J. E. Fitzgerald and members of the Clark lab for helpful conversations. DAC and this research were supported by NIH R01EY026555, NIH P30EY026878, NSF IOS1558103, a Searle Scholar Award, a Sloan Fellowship in Neuroscience, the Smith Family Foundation, and the E. Matilda Ziegler Foundation.

## Appendix A Visual stimuli used in simulations and imaging experiments

In this appendix, we describe in detail all stimuli used in this work.

### ON and OFF edges (Figure 1)

ON and OFF edges were constructed by placing white (respectively black) edges on a gray background. All edges translated at 30°/s.

### Sinusoid grating stimuli (Figures 1 and 2)

Sinusoid grating stimuli were constructed as in (Badwan et al., 2019; Creamer et al., 2018; Maisak et al., 2013). Briefly, rightward- and leftward-drifting gratings were constructed as

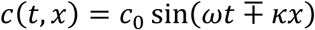

where *c*_0_ is the input contrast, *ω* is the temporal frequency in units of radians per second, *κ* is the spatial frequency in units of radians per degree, and the negative sign is taken for rightward-drifting gratings. To assess whether our model is temporal-frequency-tuned, we computed the fraction of the total variance in a spatiotemporal frequency sweep of its responses accounted for by a separable approximation resulting from its singular value decomposition (Creamer et al., 2018). Counterphase gratings were constructed as

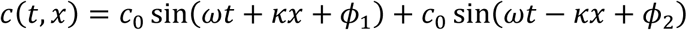

where *ϕ*_1_ and *ϕ*_2_ are uniformly sampled phase offsets, over which we average in all analyses. Gratings containing preferred- and orthogonal-direction motion were constructed as

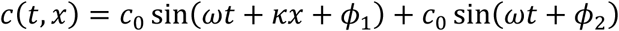

The linearity analysis in Figure 2A was applied to T5 cells (Wienecke et al., 2018), following a previously developed protocol (Jagadeesh et al., 1993). This analysis relies upon the fact that a drifting sinusoid grating may be decomposed into a sum of counterphase gratings as

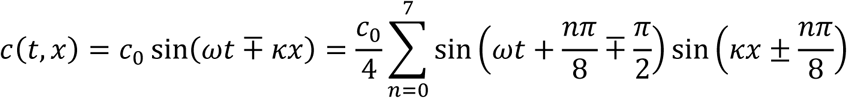

Therefore, if a system is linear, its scaled, summed response of a linear system to counterphase gratings with these phase shifts will be equal to its response to the corresponding drifting grating. By comparing the linear prediction of the drifting grating response to the actual response, one may assess a system’s linearity.

To assess the orientation- and directional-tuning of the model with sinusoid gratings in Figure 2D, we defined a two-dimensional grating

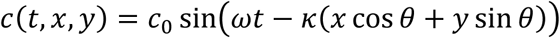

where the angle *θ* defines its orientation. In this analysis, we assume that the ring of detectors is located at *y* = 0, and that the Gaussian spatial filter is symmetric in *x* and *y*. Static gratings were formed by setting *ω* = 0. We note that our convention for the orientation of a static grating differs from the original manuscript (Fisher et al., 2015); we define the orientation as the angle between the normal to the apparent edge and the preferred direction rather than the angular position of the edge itself. Therefore, in our convention the preferred orientations and directions align.

### Apparent motion stimuli (Figure 3)

Single-bar stimuli were constructed as previously published (Gruntman et al., 2018; Salazar-Gatzimas et al., 2018). Briefly, 5° (respectively 2°) black or white bars were placed on a gray background, and presented for one second (respectively 160 ms) to match (Salazar-Gatzimas et al., 2018) (respectively (Gruntman et al., 2018)). Bar pair apparent motion stimuli were constructed as in (Salazar-Gatzimas et al., 2018). Briefly, 5° black or white bars were placed on a gray background and presented for one second. To create apparent motion, a second black or white bar was added 150 ms after the onset of the first bar at a neighboring spatial location. Responses to these bar pair apparent motion stimuli were aligned such that the location of the lagging bar matched the location of peak single-bar responses, as in (Salazar-Gatzimas et al., 2018). Flashed apparent motion stimuli were constructed similarly to those presented to T4 and T5 cells in (Gruntman et al., 2018; Haag et al., 2016). Briefly, 4.5° white bars were placed on a gray background and were presented for 400 ms in sequential spatial positions.

### Noise stimuli and linear receptive field extraction (Figure 4)

As previously published (Salazar-Gatzimas et al., 2016), we extracted linear receptive fields from responses to uncorrelated binary stimuli composed of 5° black or white bars, updated at 60 Hz. We estimated the linear receptive field from these responses using reverse correlation (Chichilnisky, 2001). Ternary noise stimuli with pairwise correlations were constructed as in (Salazar-Gatzimas et al., 2016). Briefly, the contrast of the correlated noise stimulus was given as

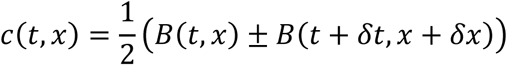

where *B*(*t*, *x*) is an uncorrelated binary stimulus composed of 5° black or white bars, and addition (respectively subtraction) generates positive (respectively negative) correlations. The stimulus was updated at a fixed rate, and the temporal offset *δt* was taken to be one cycle, with its sign determining whether the stimulus was oriented in the preferred or null direction. The spatial offset *δx* was fixed to be one bar width. As shown in (Salazar-Gatzimas et al., 2016), the autocorrelation function of this stimulus, with spacetime discretized by the bar width and sampling rate, is

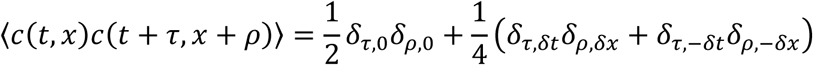

where *δ*_*i,j*_ is the Kronecker delta.

### Three-point glider stimuli (Figure 4)

As in previous studies (Clark et al., 2014; Fitzgerald and Clark, 2015), we constructed three-point glider stimuli following (Hu and Victor, 2010). Briefly, these binary stimuli enforce correlations over space and time among triplets of pixels. Three-point gliders may be categorized into four types: converging gliders with positive parity (con+), converging gliders with negative parity (con-), diverging gliders with positive parity (div+), and diverging gliders with negative parity (div-). Letting *ρ* be the pixel spacing and *δ* be the frame duration (the inverse of the update rate), the update rules for each of the four three-point glider types are (see kymographs in **Figure 4**):

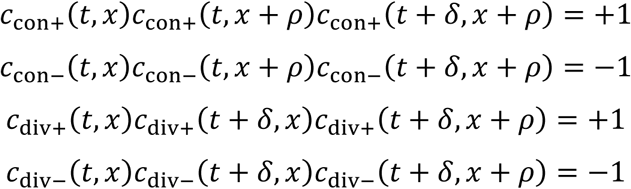

The direction of the displacement of the spatial mean location of each triplet is inverted by inverting the sign of the pixel spacing. Starting from an initial seed state, the values of each pixel at each timepoint are determined by these update rules using the surrounding pixels’ values. As we simulate the full 360° of visual space, we use periodic boundary conditions to avoid undetermined edge pixel values. As in previous studies (Clark et al., 2014; Fitzgerald and Clark, 2015; Leonhardt et al., 2016), the pixel spacing was taken to be 5° in both imaging experiments and numerical simulations. In imaging experiments, visual stimuli were generated and presented as described in previous studies (Badwan et al., 2019).

### Random checkerboard stimuli (Figure 4)

Random checkerboard stimuli were constructed as in (Badwan et al., 2019). Briefly, 5° black or white bars were placed at random with a density of 40% on a gray background. The resulting checkerboards were then rigidly translated at a velocity of 100°/s. When combining rightward- and leftward-moving stimuli, summation was defined such that two white bars summed to white, two black bars summed to black, and one white and one black bar summed to gray. Therefore, the contrast of the composite stimulus matched that of the individual components, though its density rose to 64%.

### Natural scene stimuli (Figure 5)

Following prior work (Chen et al., 2019; Clark et al., 2014; Fitzgerald and Clark, 2015), we generated a left-right symmetric ensemble of natural scenes by drawing independent row and column samples from the database gathered by (Meyer et al., 2014). In this ensemble, scenes were rigidly-translated at velocities sampled from a Gaussian distribution with a standard deviation of 100°/s, which roughly matches typical rotational velocities of walking flies (DeAngelis et al., 2019; Katsov and Clandinin, 2008). To convert the scenes to contrast signals, we spatially filtered each image with the photoreceptor kernel to generate blurred images *I*_blur_, and then used a Gaussian kernel with a standard deviation of 20° to estimate locally-averaged images *I*_mean_ (Chen et al., 2019). The contrast signal was then defined as

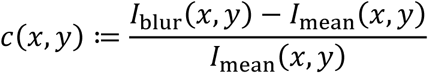

As in previous studies of coactivation (Salazar-Gatzimas et al., 2018), the coactivations in Figure 5C were computed as normalized inner products of response timeseries. For all analyses in Figure 5, we used an ensemble with 10^6^ elements.

## Appendix B Parameter value selection

In this appendix, we briefly describe how we selected values of the weighting parameters *g*_exc_/*g*_leak_ and *g*_inh_/*g*_leak_. We evaluated the model solely based on its ability to produce direction-opponent average responses to 1 Hz, 45° sinusoid gratings similar to those measured in T4 cells (Badwan et al., 2019). To do so, we considered the direction selectivity index and analogous indices of direction-opponency and orthogonal direction enhancement, defined as

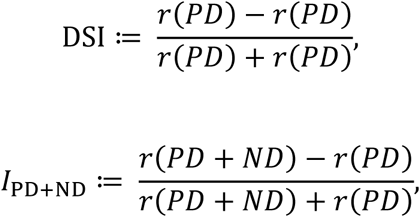

and

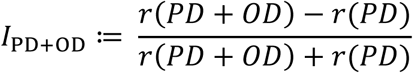

As shown in **Figure B1**, there exists a broad region of parameter space for which the model produces responses with a similar degree of direction-opponency to that measured in T4 cells without significant PD+OD enhancement. We therefore made a simple choice of round-number values within that region.

**Appendix Figure B1:**
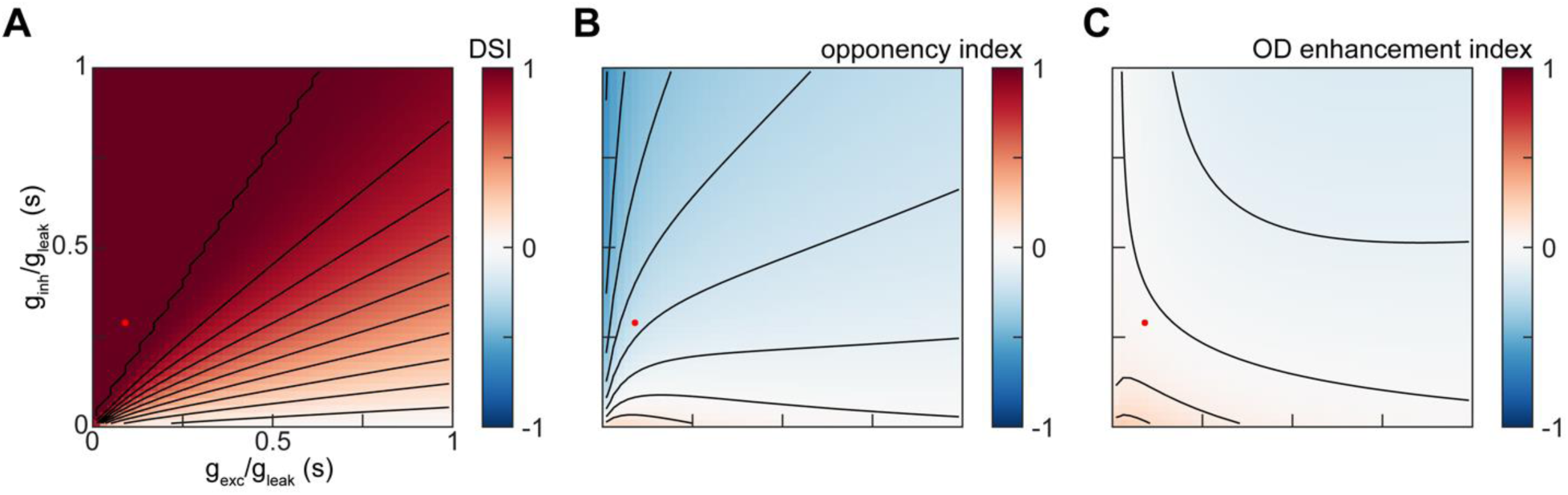
Sweeping the parameters of the T4 synaptic model. A. The direction-selectivity index of the T4 synaptic model’s responses to drifting sinusoid gratings as a function of *g*_exc_/*g*_leak_ and *g*_inh_/*g*_leak_. Red dot indicates the selected values of *g*_exc_/*g*_leak_ = 0.1 and *g*_inh_/*g*_leak_ = 0.3. B. As in (A), but for the opponency index. C. As in (A), but for the OD enhancement index.

## Appendix C LNLN cascade factorization of the T4 synaptic model

In this appendix, we show how our T4 synaptic model may be factorized as a product of linear-nonlinear-linear-nonlinear (LNLN) cascades representing the numerator and denominator of the biophysical nonlinearity. The response *C* of the full model at each point in spacetime is given in terms of the filtered contrast signal *s* as

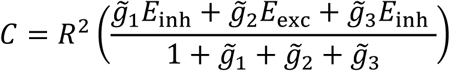

where we have defined 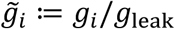 for brevity. Noting that the denominator of this expression is always positive, we may re-express the response as

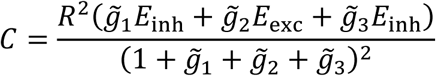

hence the full EMD model admits a factorization into a product of LNLN models as

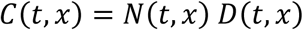

where

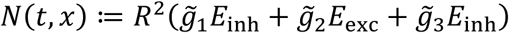

and

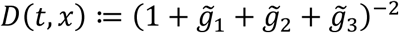

which is bounded as *D*(*t*, *x*) ≤ 1. Because *D*(*t*, *x*) ≤ 1, *C*(*t*, *x*) ≤ *N*(*t*, *x*).

The denominator LNLN cascade *D* is the result of applying a convex function (*x*^−2^ for *x* > 0) to a non-negative linear combination of LN models with convex nonlinearities. Therefore, it cannot generate direction-opponent (DO) average responses to sinusoid gratings. The proof of this proposition is a minor extension of our previous results on LNLN models with continuously-differentiable convex nonlinearities and non-negative secondary linear filters (Badwan et al., 2019). We define the soft ramp function

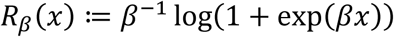

which is a continuously differentiable, monotone increasing, non-negative, and convex function of *x* for all positive *β*. As *β* → ∞, *R*_*β*_(*x*) → *R*(*x*) pointwise. By continuity, defining *D*_*β*_ (*t*, *x*) using *R*_*β*_, we have 0 ≤ *D*_*β*_(*t*, *x*) → *D*(*t*, *x*) ≤ 1 as *β* → ∞. We denote the nonlinear functional corresponding to the spacetime average of *D*_*β*_(*t*, *x*) for some input stimulus *f* as *D*_*β*_ [*f*]. As we have the integrable constant dominating function 1, by the Lebesgue dominated convergence theorem we have 0 ≤ *D*_*β*_[*f*] → *D*[*f*] ≤ 1 as *β* → ∞ (Stein and Shakarchi, 2009). By the result of (Badwan et al., 2019), we know that *D*_*β*_ [*PD* + *ND*] ≥ *D*_*β*_[*PD*] and *D*_*β*_[*PD* + *ND*] ≥ *D*_*β*_[*ND*], where *D*_*β*_ [*PD*], *D*_*β*_[*ND*], and *D*_*β*_ [*PD* + *ND*] are the average responses to PD, ND, and PD+ND sinusoid gratings, respectively. As these inequalities hold pointwise for all positive *β*, by taking *β* → ∞ we may obtain *D*[*PD* + *ND*] ≥ *D*[*PD*] and *D*[*PD* + *ND*] ≥ *D*[*ND*]. Therefore, the denominator LNLN cascade cannot generate DO average responses to sinusoid gratings.

However, as the numerator LNLN model is the result of applying a convex function to a non-convex linear combination of LN models with convex nonlinearities, we cannot analytically exclude the possibility that it could generate DO average responses to sinusoid gratings using the results of (Badwan et al., 2019). In fact, numerical simulation shows that it can generate DO average responses to sinusoid gratings, though it generates strong PD+OD enhancement (**Figure C1**). It also generates DO responses over a smaller region in spatiotemporal frequency space than the full model. If one replaced the infinitely sharp ramp functions with more biophysically plausible soft rectifiers, the numerator LNLN cascade would be well-approximated for small input contrasts by a LN model with a quadratic nonlinearity. Therefore, it could not generate DO average responses for sufficiently small input contrasts. However, even in the limit in which both the numerator and denominator are represented as LN models with quadratic nonlinearities, the full model could likely generate DO average responses. In particular, this limiting construction would resemble a type of adaptive gain model which was previously shown to generate DO average responses (Badwan et al., 2019).

**Appendix Figure C1:**
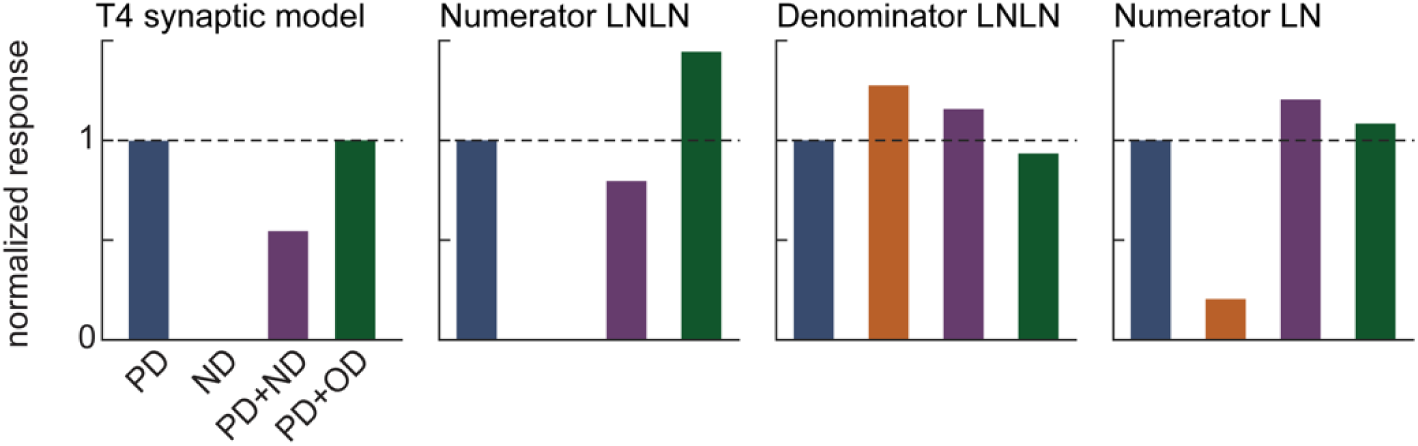
Sinusoid grating responses of different components of the LNLN factorization. *From left to right:* Average responses of the full T4 synaptic model, the numerator LNLN cascade, the denominator LNLN cascade, and the numerator LN cascade to 1 Hz, 45° sinusoid gratings. All responses are normalized by the response of the given component to a grating drifting in the PD of the full model.

